# Variation in *Plasmodium falciparum* sexual commitment rates and responses to environmental modification

**DOI:** 10.1101/2022.02.14.480314

**Authors:** Lindsay B. Stewart, Aline Freville, Till S. Voss, David A. Baker, Gordon A. Awandare, David J. Conway

**Author notes:** These authors contributed equally to the research (Lindsay B. Stewart, Aline Freville).

## Abstract

Asexual blood stage malaria parasites must produce sexual progeny in order to infect mosquitoes. It is therefore important to understand the scope of variation in sexual commitment rates of the most important human parasite *Plasmodium falciparum*, and to assess responses to conditions that may affect this commitment. First, using a genetically modified line with inducible elevated sexual commitment we compared two different methods of measuring the rates, either staining the Pfs16 marker in very early gametocytes or counting gametocytes after further development. The methods showed correlated estimates in evaluation across multiple independent replicate assays, with higher sensitivity and precision being achieved by Pfs16 marker detection in early gametocytes, so this method was used to survey a wide range of *P. falciparum* lines. A panel of six recently culture-established clinical isolates from Ghana showed mean sexual commitment rates per cycle ranging from 3.3% to 12.2%, with some significant differences among isolates as well as variation among biological replicates of each. Comparing 13 long-term laboratory-adapted lines of diverse origins, eight had sexual commitment rates similar to those of the recent clinical isolates, with means across multiple biological replicates for each line ranging from 4.7% to 14.5%, while four lines had significantly lower rates with means ranging from 0.3 to 1.6%, and one line with a non-functional *ap2-g* gene always showed zero sexual commitment. Among a subset of parasite lines, adding choline to suppress commitment had quantitatively variable effects, although these were significant in most assays of lines that had relatively high rates. This study indicates the importance of multiple assay replicates and comparisons of diverse isolates to understand natural and culture-induced variation in this key reproductive trait, relevant to investigating potential effects of parasite adaptation on transmission.

**Author Summary:** Only sexual malaria parasites are transmitted from humans to mosquitoes, so it is vital to understand variation in sexual commitment rates of blood stage malaria parasites, and responses to conditions that affect this. We compared two different methods of measuring the rates in a *Plasmodium falciparum* line with engineered sexual commitment variation, demonstrating higher sensitivity and precision by detection of an early differentiation marker, and this method was then used to survey diverse *P. falciparum* lines in culture with extensive biological replicate measurements. Recent clinical isolates from Ghana showed mean sexual commitment rates per cycle ranging from 3% to 12%, with significant differences among isolates as well as variation among biological replicates. Long-term laboratory-adapted lines of diverse origins had a wide range, most having rates similar to those of the clinical isolates, while a minority consistently had much lower or zero rates. There was quantitative variation among lines in the effects of adding choline to suppress commitment, although most assays of lines that had relatively high rates showed significant effects. Performing multiple biological replicates and comparisons of recent parasite isolates is vital to understand intra-specific variation in this important reproductive trait, and move towards investigating direct effects on disease transmission.

## Introduction

For malaria to be transmitted, asexual blood stage parasites must undergo differentiation which leads to formation of male and female gametocytes that are required to infect mosquitoes. This process involves an initial parasite commitment phase, when the switch to sexual development is induced, followed by a longer developmental phase to produce the mature gametocyte forms. From an evolutionary or ecological perspective, it is important to understand what determines the rate of commitment to sexual forms [1], and how this may vary within a species [2]. It is particularly vital for understanding *P. falciparum* transmission as more effective control is needed to reduce the global malaria burden [3], which is particularly challenging as chronic asymptomatic infections are often highly infective to mosquitoes [4].

Research on malaria parasite sexual conversion has been long established [5], and progress has been made in elucidating key aspects of the mechanism. An ApiAP2-family transcription factor AP2-G is essential for sexual conversion, controlling expression of genes critical to early gametocytogenesis [6-9]. The *ap2-g* locus is clonally variably expressed, with histone 2 lysine 9 trimethylation (H3K9me3) marks of heterochromatin silencing initially indicating that its expression could be epigenetically regulated [10], and conditional depletion of heterochromatin protein 1 (HP1) from the promoter was shown to increase *ap2-g* transcription and sexual conversion [11]. Furthermore, in *P. falciparum* a gametocyte development protein GDV1, that was previously linked with sexual commitment [12], is involved in removal of HP1 from the *ap2-g* locus so that over-expression of *gdv1* causes elevated sexual commitment rates [13].

The classic model is that sexual commitment occurs in the intra-erythrocytic cycle prior to the cycle in which parasite sexual differentiation occurs, based on a finding that in static culture individual schizonts of the 3D7 parasite clone yield spatially-clustered intra-erythrocytic progeny that tend to differentiate similarly, either sexually or asexually [14]. However, there is recent evidence that some sexual commitment may also occur in the same cycle as sexual differentiation, if *ap2-g* expression is induced to start very early after erythrocyte invasion during the ring stage [15], although it is not known how commonly this occurs in parasites under normal conditions of infection or laboratory culture. It is difficult to study *P. falciparum* gametocyte conversion rates precisely in natural infections, as most stages of developing gametocytes do not occur in the peripheral circulation and are therefore not detected in blood samples, although circulating ring stage parasites that are sexually committed may become detectable by developing assays for early markers such as AP2-G or GEXP02 [16, 17]. Correlations between *P. falciparum* gametocyte numbers and other measured variables in human hosts, such as age or hemoglobin levels, may not be causally related to varying gametocyte conversion rates and are not well replicated across different studies [18]. Experiments with other malaria parasite species in laboratory mice indicate sexual commitment to be a plastic phenotype that can respond to the *in vivo* environment [19]. There are few data relating to *P. falciparum* gametocyte conversion in experimentally induced human infections, but modelling of results from human experimental infections with the parasite clone 3D7 has indicated that the conversion rate per cycle is usually less than 1% [20], and older data on induced malaria infections with strains that no longer exist suggest broadly similar average rates [21].

Early studies of *P. falciparum* in culture indicated that gametocyte production varied among parasite clones [22, 23], and also varied with culture conditions, although precise estimates of conversion rates per cycle were not established. Subsequent studies estimated gametocyte conversion rates of several laboratory-adapted *P. falciparum* clones to be usually above 1% per cycle when grown at low parasitaemia [6, 24], with higher rates if parasites were stressed at high parasitaemia or by antimalarial drug treatment [14, 24].Parasites appear to respond to metabolic challenges during growth, although most cellular mechanisms remain to be identified [25]. An increase in gametocytogenesis may be triggered by cellular stress involving redox perturbation [26, 27], by adding low doses of the antimalarial artemisinin at the early trophozoite growth stage [28], or by adding parasite-conditioned culture medium [29] or extracellular vesicles from such medium [30]. Notably, gametocyte conversion rates in cultured lines are suppressed by lysophosphatidylcholine (LysoPC) within serum, or by adding choline as a supplement in serum-free medium [31, 32]. In a study of parasite development during the initial *ex vivo* cycle cultured from clinical samples in Ghana, the proportions of parasites showing conversion to gametocytes correlated positively with level of parasitaemia and negatively with concentration of LysoPC in plasma of patients, supporting a hypothesis that developmental responses to metabolic stress may occur during natural infections [33].

Despite these advances, the quantitative scope of sexual commitment rate variation in *P. falciparum* has not yet been clearly defined. Estimates of commitment rates have previously varied due to methodological differences or to biological differences between nominally similar parasite lines, and there are few controlled comparisons of different isolates employing biological replicate measurements. Performing multiple independent replicate assays is important, because some individual lines may have temporally variable phenotypes and because precision is generally increased with repeated measurements. In this study, we first compared different methods of measuring sexual commitment rates, and then employed the method with greater quantitative sensitivity and accuracy to conduct multiple biological replicate assays on recently adapted clinical isolates from Ghana as well as a broad selection of long-term laboratory adapted lines representing most of those that are commonly cultured globally. Although each clinical isolate showed variability among repeated assays, overall there are significant differences among isolates, indicating potentially adaptive variation in sexual commitment within this important parasite species. Separate assays of selected lines on the suppressive effect on sexual commitment by adding choline to serum-free culture media identified significant quantitative variation among lines, which further encourages research to test for relevance to disease transmission.

## Results

### Comparison of sensitivity and precision of two different assays to measure gametocyte conversion rates

Two different assay methods were compared to quantify sexual commitment rates in multiple independent experiments, using a genetically engineered *P. falciparum* clone (3D7/iGP_D9) to over-express GDV1 with a destabilising domain (DD) in the presence of the stabilising reagent Shield-1 which leads to elevated gametocyte conversion [34]. Method 1 involved differential counting of stage I gametocytes detected by anti-Pfs16 immunofluorescence staining as a proportion of all parasites developing beyond ring stages in the next cycle, while Method 2 involved Giemsa-stained slide microscopy to compare the next cycle ring stage parasitaemia (referred to as day 0 in this assay, D0) with the parasitaemia of morphologically-defined stage II gametocytes (at day 4, D4) (details given in the Materials and Methods section). In each of multiple biological replicate experiments, both methods clearly show that the gametocyte conversion rate is very significantly higher when GDV1 is overexpressed and stabilised with Shield-1 (GDV1-DD-ON) compared to when GDV1 overexpression is disabled by the absence of Shield-1 (GDV1-DD-OFF) (Figure 1, Supplementary Table S1).

**Figure 1.**
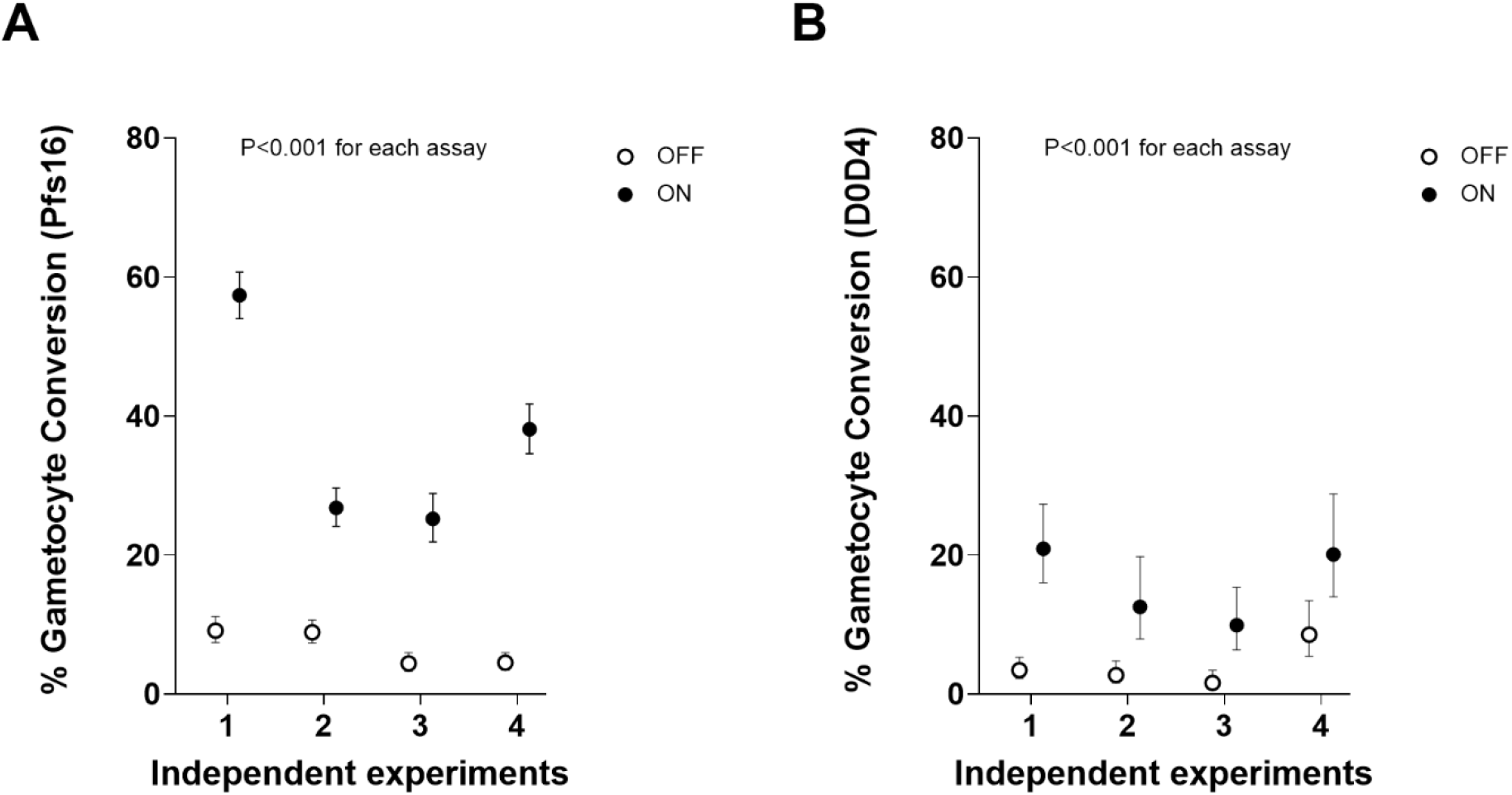
Comparison of the sensitivity and precision of two different methods for measuring *P. falciparum* sexual commitment in terms of gametocyte conversion rates. Multiple paired replicate assays were performed with the parasite clone 3D7/iGP_D9 [34] which allows inducible overexpression of GDV1 fused to a destabilization domain (DD) stabilized in the presence of Shield-1 reagent to provide controlled elevation of sexual commitment (closed circles indicate GDV1–ON, open circles GDV1-OFF). Both methods show the expected significant elevation in gametocyte conversion in the presence of Shield-1 reagent in all individual assays (asterisks indicate P<0.0001 for each of the individual replicate assays). In each panel, the green dotted line denotes the mean for GDV1-ON and the red dotted line for GDV1-OFF. **A**. The gametocyte conversion rate (with 95% CI) for each of seven independent experiments using Method 1 involving staining of the gametocyte marker Pfs16, yielding a mean conversion rate of 38.3 % under GDV1-ON and 6.3% under GDV1-OFF conditions **B**. The gametocyte conversion rate (with 95% CI) for each of four experiments using Method 2 which involves a comparison of Day 4 (D4) gametocyte parasitaemia with D0 ring-stage parasitaemia (assays paired with the first four independent experiments shown for Method 1), yielding a mean conversion rate of 15.9 % under GDV1-ON and 4.1 % under GDV1-OFF conditions. Exact parasite and erythrocyte counts and statistical analyses for all assays are detailed in Supplementary Table S1.

In all biological replicate assays, Method 1 showed a marked difference in gametocyte conversion rate between the GDV1-DD ON and OFF conditions, with a mean conversion rate of 36.9 % under ON and 6.7 % under OFF conditions (Figure 1). In comparison, Method 2 yielded an estimated mean conversion rate of 15.9 % under ON and 4.1 % under OFF conditions, with the rates for Method 2 being lower than for Method 1 in all replicates for which both were tested in parallel. The precision of the estimates for each of the biological replicates was higher for Method 1, with relatively narrow confidence intervals for each experimental replicate as this method is based on direct counts of developmentally discrete parasites. In comparison, Method 2 yielded wider confidence intervals as it is based on a ratio-of-ratio proportion of two temporally sequential counts of parasitaemia which involves more statistical sampling error (Supplementary Table S1), and this method also depends on successful parasite development over a longer time in culture.

### Correlation between gametocyte conversion rate estimates from different assay methods

To test for overall correlation between the estimates from the two assay methods, analysis was performed on data from a broader range of paired assays. In addition to the eight measurements from paired assays performed in the first comparison (Figure 1) nine additional measurements of the 3D7/iGP_D9 clone were performed with different concentrations of the Shield-1 reagent, to yield a total of 17 paired measurements with both methods for analysis (Supplementary Table S2). Overall, this showed a high and significant nonparametric rank correlation between the results from each assay method (Spearman’s rho = 0.70, P = 0.003), with Method 1 generally giving higher conversion rate estimates (Figure 2) with narrower 95% confidence intervals (Supplementary Table S2). While this supports both assays as separately giving informative relative estimates of gametocyte conversion, Method 1 was chosen for subsequent quantitative analyses of additional lines and culture conditions given its significantly higher sensitivity and tighter precision (Figures 1 and 2, Supplementary Tables S1 and S2).

**Figure 2.**
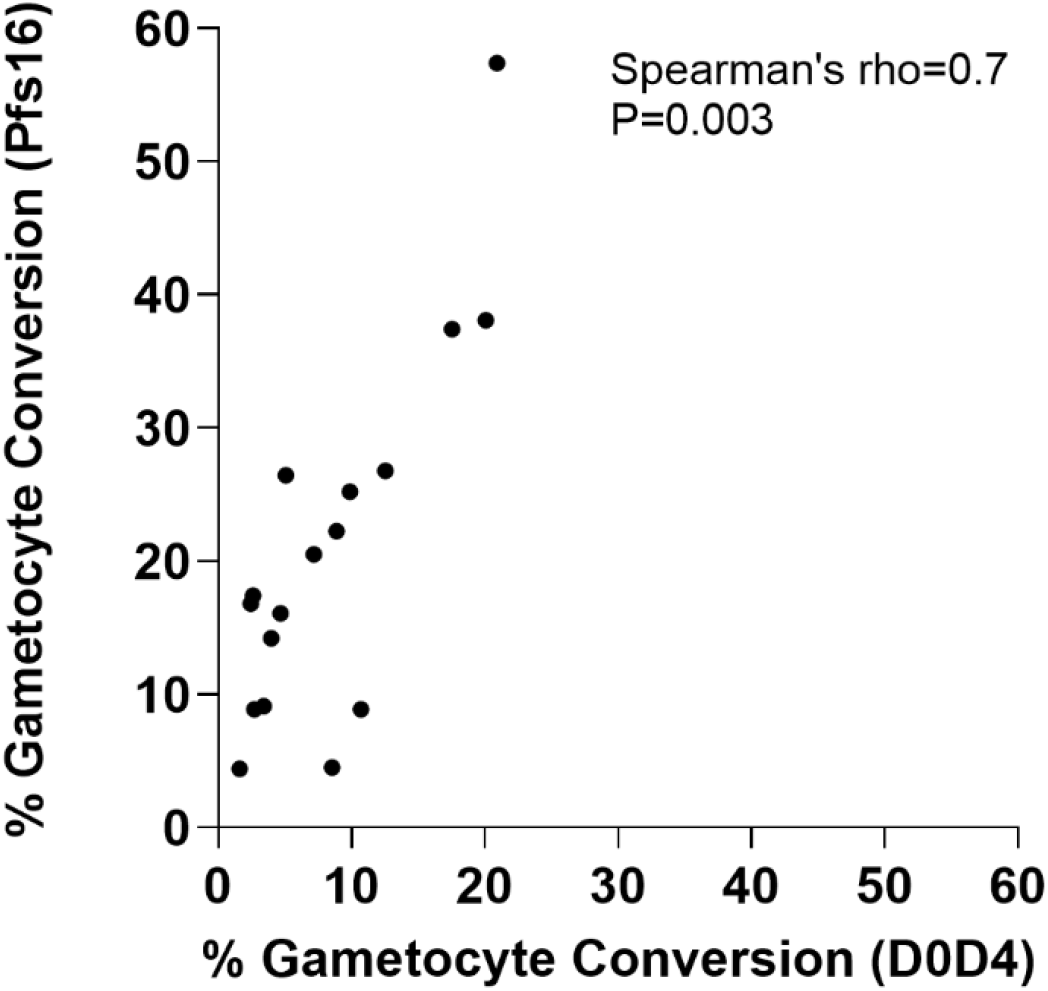
Significant correlation in the estimates of gametocyte conversion rate obtained by two different assay methods. Paired assays were performed using Method 1 (involving Pfs16 staining to discriminate sexually from asexually-developing parasites) and Method 2 (D0D4, involving counts of parasites 4 days apart) on multiple biological replicates of 3D7/iGP_D9 with a range of concentrations of Shield-1 reagent (N=17 paired assays in total, details in Supplementary Table S2). This highly significant correlation confirms that either method gives valid measurements if used alone, whereas Method 1 is more sensitive (as indicted here and in Figure 1). Method 1 is also more precise as it involves calculation of a single numerical ratio, rather than a ratio-of-ratios based on two different counts performed 4 days apart which is highly vulnerable to effects of small numbers in either count. For visual clarity, the individual assay 95% confidence intervals are not shown on the plot, but these and numerical data of all assays are given in Supplementary Table S2.

### Variation in sexual commitment rates of recently culture-established Ghanaian clinical isolates

Recent culture-established clinical isolates from a highly endemic area in Ghana were assayed to measure gametocyte conversion rates. Six isolates were selected from a larger panel for analysis as they represent unrelated genetically unmixed infections [35], so that biological replicate assays correspond to a single parasite genotype in each case, and after culture for several months they did not have any known loss of function mutations detected in their genome sequences that would affect sexual commitment [35]. These isolates show mean gametocyte conversion rates that range from 3.3% to 12.2% (Figure 3, Supplementary Table S3), and there were significant differences in pairwise comparisons (Mann-Whitney tests on results of a minimum of six biological replicate assays performed for each isolate, Supplementary Table S4). For example, isolate 296 had a significantly higher conversion rate than three of the others (isolates 272, 289 and 292), and isolate 289 had a significantly lower rate than three of the others (292, 293 and 296)(Supplementary Table S4).

**Figure 3.**
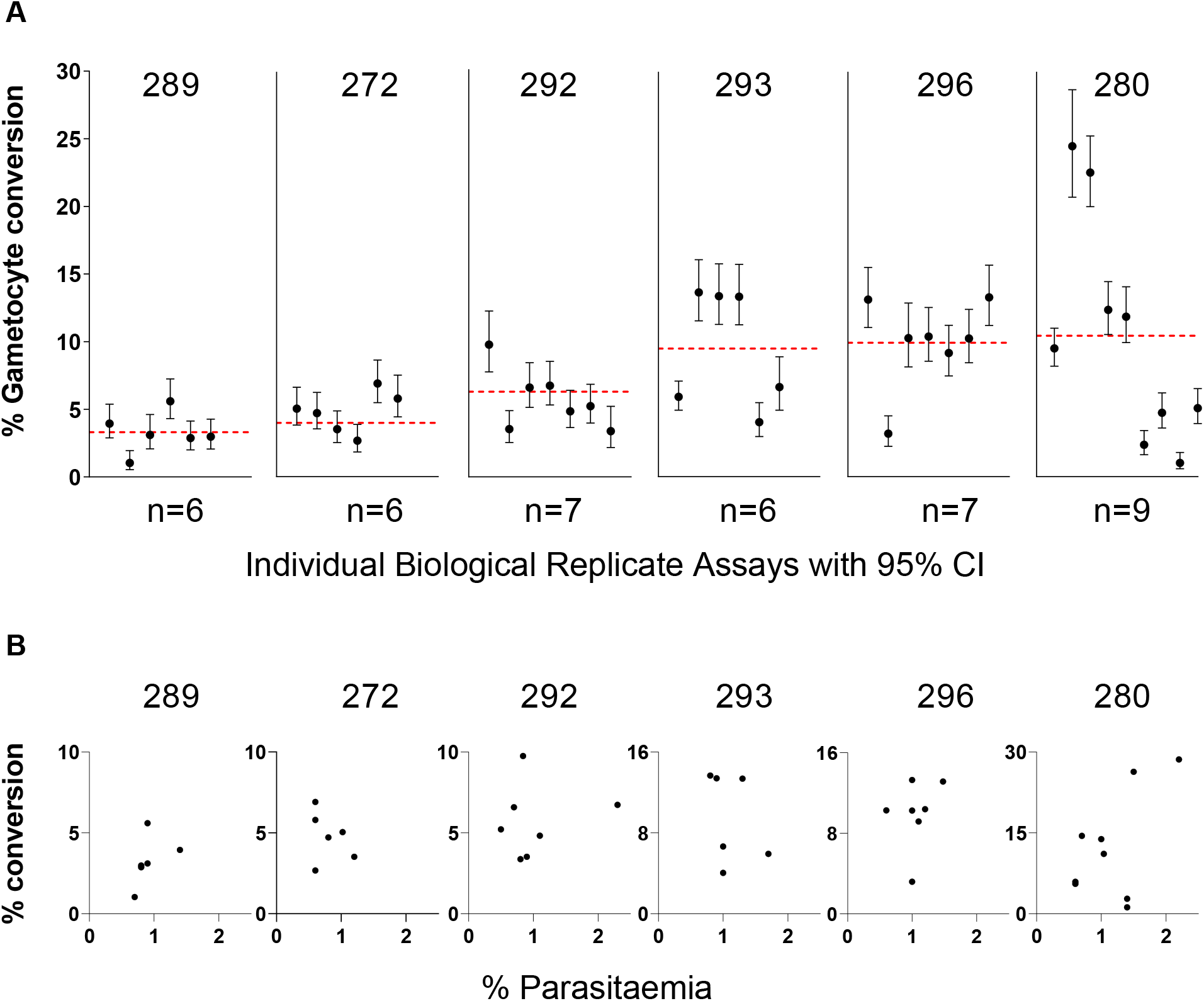
Gametocyte conversion rates of six recently culture-established Ghanaian clinical isolates. These were selected for study as they are unrelated single genotype infections, chosen from a larger number of samples from a highly endemic population that contained many multiple genotype infections [35]. **A**. For each isolate, a minimum of six biological replicate assays was performed, and the estimate from each replicate with 95% confidence intervals (based on numbers of parasites counted) is plotted, while the red dotted line represents the mean across replicate assays. There were significant differences in pairwise comparisons (Mann-Whitney tests, Supplementary Table S4), isolate 296 having a significantly higher conversion rate than three others (isolates 272, 289 and 292), and isolate 289 having a significantly lower rate than three others (292, 293 and 296)(Supplementary Table S4). There were no significant correlations with length of time in culture, or batches of erythrocytes used for the different replicate assays (these details and all numerical data values are given in Supplementary Table S3). **B**. For each isolate the gametocyte conversion rate of individual replicates is plotted in relation to the culture parasitaemia in the previous cycle (between 0.5 and 2.5% for all experimental replicates). 95% confidence intervals individual conversion rate points are not shown as they are the same as in panel A. There was no systematic trend towards positive or negative correlation, except a marginally significant positive correlation for isolate 289 (P = 0.03) which may indicate slight enhancement at higher parasitaemia or a chance finding due to multiple comparisons.

There was also significant variation in gametocyte conversion rates among individual biological replicate assays of each isolate, with non-overlapping 95% confidence intervals of some replicates (Figure 3A). The variation among biological replicate assays for each line was not correlated with the culture parasitaemia, which was always between 0.5 and 2.5% (Figure 3B and Supplementary Table S3). Overall, there was no significant evidence of reduction in gametocyte conversion rates over time in culture (Supplementary Table S3), and variation among replicates did not correspond with the use of different erythrocyte batches for culture (Supplementary Table S3).

### Range of gametocyte conversion rates among diverse long-term cultured laboratory lines

To compare with these recent clinical isolates, a panel of 13 long-term culture-adapted *P. falciparum* lines of diverse origins was tested, with multiple biological replicate assays (Figure 4, Supplementary Table S5). The clone F12 (containing a nonsense mutation in *ap2-g*)[6] here showed zero conversion in all replicate assays (Figure 4A, Supplementary Table S5). Four of the other lines showed consistently low levels of gametocyte conversion, with means ranging between 0.3% and 1.6%, significantly lower than the conversion rates of the remaining lines (Mann-Whitney tests on pairwise comparisons, Supplementary Table S6). The lines with the lowest conversion rates (D10 and T9/96) were previously described as having multiple gene deletions in a sub-telomeric region of chromosome 9 including the *gdv1* gene [36]. The 3D7 line may contain undetected mutants within bulk culture, as 3D7 has been elsewhere shown to be a source of parasite subclones with loss of function mutations (including the *ap2-g* nonsense mutation in parasite subclone F12 [6], deletion of *gdv1* in subclone 3D7.G_def_ [12] and truncation of *gdv1* in two other subclones [37]). The Palo Alto line was previously shown to have undergone sub-telomeric gene deletions on different chromosomes [36, 38] but not to have lost the *gdv1* locus on chromosome 9 [36, 39], although the possibility of a loss-of-function mutation has not been excluded.

**Figure 4.**
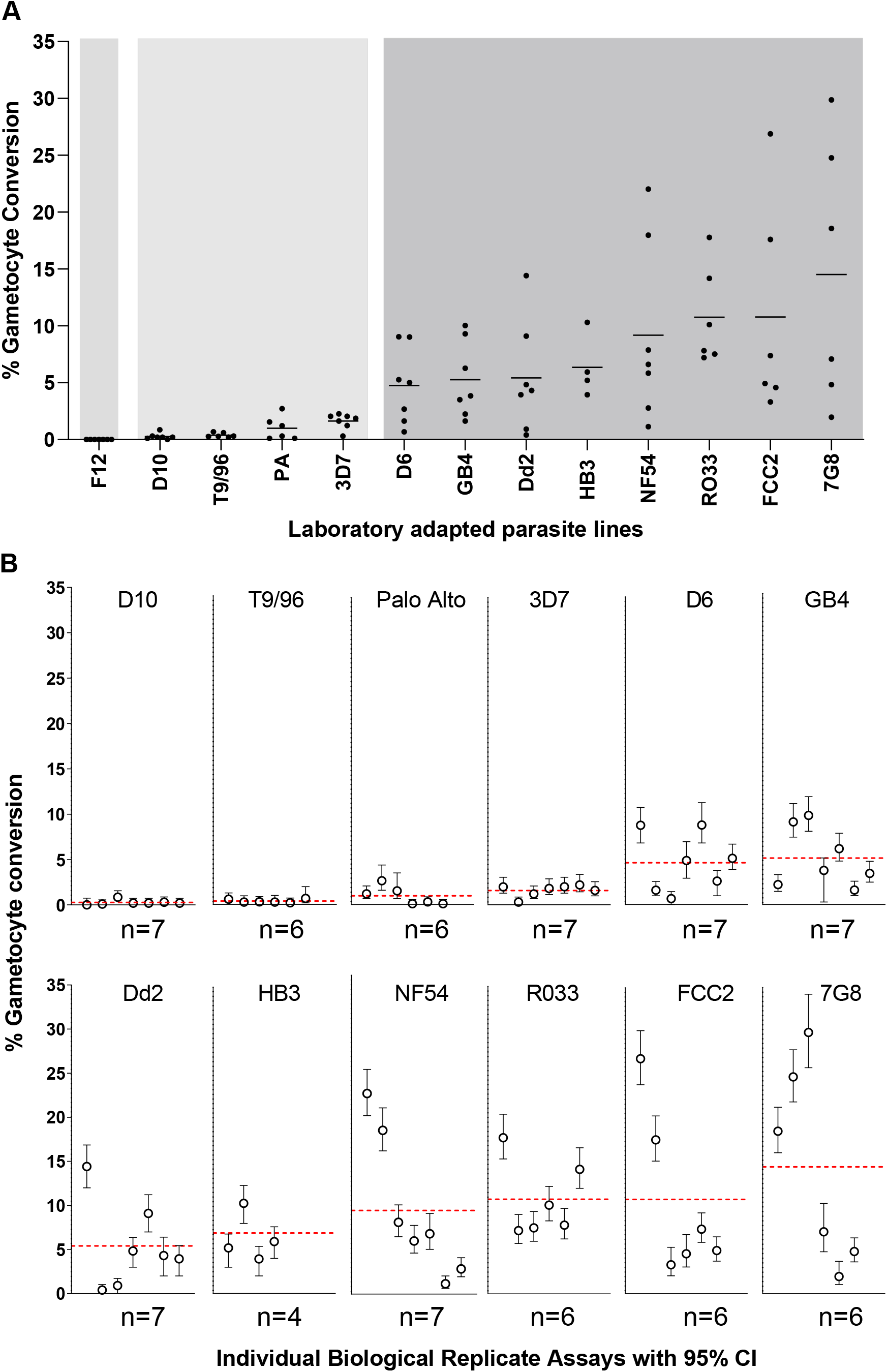
Gametocyte conversion rates of 13 long-term culture adapted *P. falciparum* lines of diverse geographical origins. Each different parasite line was tested with multiple independent biological replicate assays, most individual lines having at least six replicates. All counts and calculated values for all replicates are given in full in Supplementary Table S5. **A**. For each parasite line, the conversion rates are shown for each of the biological replicates and the horizontal line shows the mean across all biological replicates, with a spectrum of variation among the different lines, means ranging from 0% to 14.5%. The background with different shades of grey show that the conversion rates separate into three different groups, based on pairwise comparisons of all replicates between the parasite lines (Mann-Whitney tests, Supplementary Table S6): One line (F12) showed no conversion, four lines (D10, T9/96, Palo Alto, and 3D7) showed consistently low conversion (means between 0.3% and 1.6%), and eight lines (D6, GB4, Dd2, HB3, NF54, RO33, 7G8, and FCC2) showed higher levels of conversion (means between 4.7% and 14.5%). **B**. Variation in the gametocyte conversion rates among independent replicate assays for each line is shown in more detail by including 95% CI of the counts for each assay of each line, except F12 for which there was zero conversion in all assays. Identical methods were used in assaying these parasite lines as for the clinical isolates shown in Figure 3, assays being conducted in parallel under the same conditions in the same laboratory. Erythrocytes from different erythrocyte donors used at various times throughout the series of experiments did not determine the variation between assay replicates (Supplementary Table S5).

For the remaining eight long-term culture-adapted *P. falciparum* lines, the gametocyte conversion rates for each showed means ranging from 4.7% (for D6) to 14.5% (for FCC2) (Figure 4A). This range is similar to that seen for the recently culture-established clinical isolates, with no significant differences between these groups (Mann-Whitney test, P = 0.47). Similar to the clinical isolates, the individual long-term culture-adapted parasite lines also showed significant variation in gametocyte conversion rates among biological replicate assays (Figure 4A), with non-overlapping 95% confidence intervals for some replicates (Figure 4B). This shows that high and variable sexual commitment rates have been maintained in a majority of the diverse cultured *P. falciparum* lines, in contrast to a minority of lines that have markedly reduced rates or zero commitment.

### Choline in culture medium suppresses sexual commitment rate with varying levels of effect among *P. falciparum* lines

As it has been previously shown that parasite sexual commitment responds to the presence or absence of choline in culture medium [13, 28, 31], we investigated five long-term culture-adapted lines (T9/96, 3D7, Dd2, NF54 and HB3) for quantitative effects of 2 mM choline compared to no added choline, performing between six and eight biological replicate assay comparisons for each parasite line. Significant suppression of sexual commitment in the presence of choline occurred in most lines (Figure 5, Supplementary Table S7). No significant effect was seen in the *P. falciparum* clone T9/96 which has a deletion of *gdv1* and several adjacent genes deleted on Chromosome 9 [36], and which has a very low rate of sexual commitment either in the presence or absence of choline, consistent with the importance of *gdv1* in responding to environmental induction [13]. The responses of the other lines to choline varied, with statistically significant suppression in sexual commitment in most of the replicate assays for NF54 and Dd2, in half of the assays for HB3, and in a quarter of the assays for 3D7 (Figure 5). Some individual assay replicates showed more than 10 fold suppression of sexual commitment rate in the presence of choline, but most showed more moderate rates (Supplementary Table S7).

**Figure 5.**
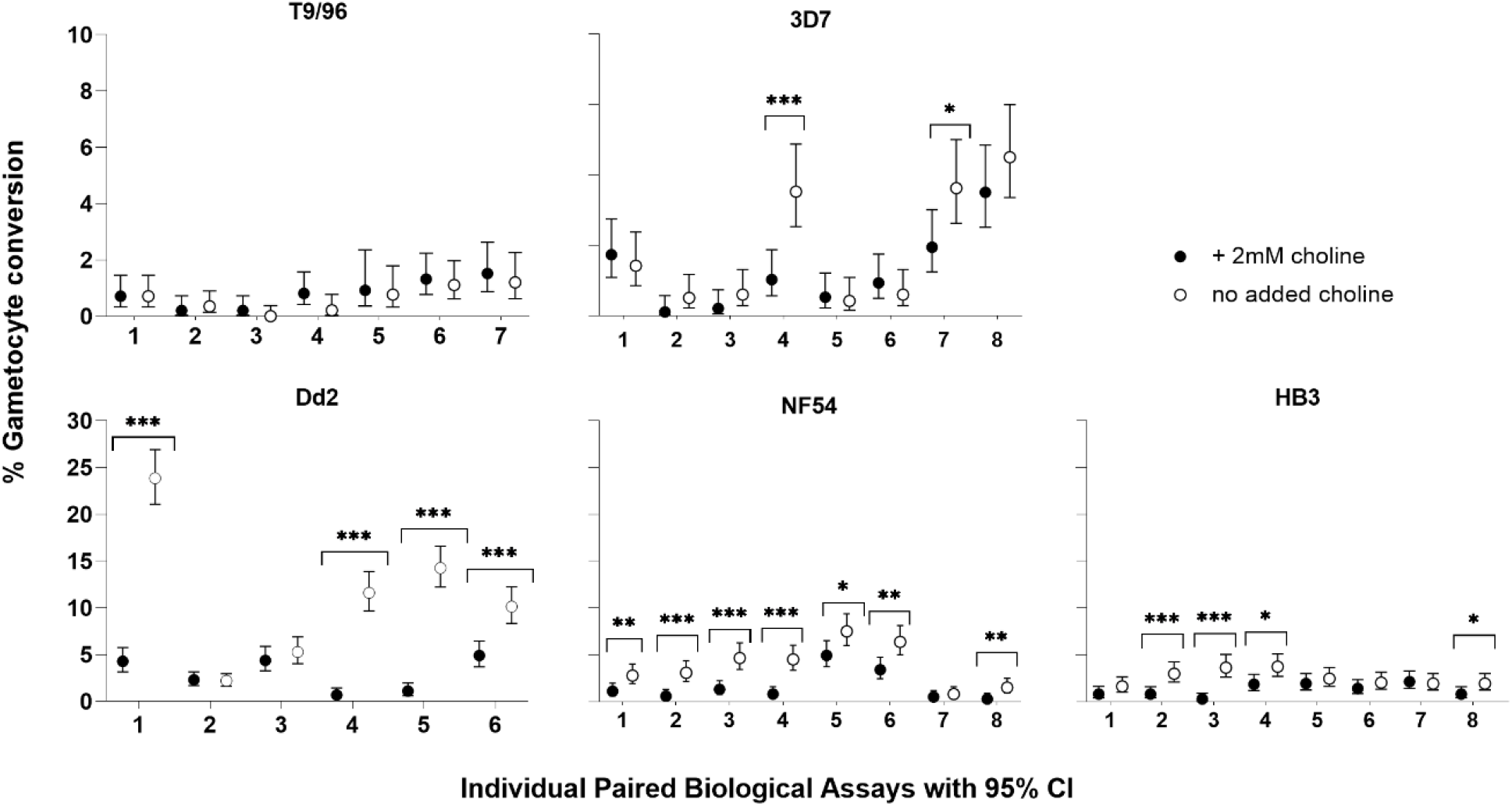
Significant variation in sensitivity of different parasite lines to choline in media to suppress sexual commitment. Closed circles represent biological replicates cultured in the presence of 2 mM choline, open circles represent biological replicates grown in the absence of choline. T9/96 shows no response to choline in any of seven biological replicate assays, whereas each of the other cultured lines show responses that vary among biological replicates. Significant differences between paired measurements with and without choline in individual replicate assays for 3D7, Dd2, HB3 and NF54 are highlighted with asterisks (Fisher’s Exact Test * P < 0.05, ** P <0.01, *** P<0.001). Across all replicates, estimated by Mantel-Haenszel rate ratios the mean effects of choline were greatest for Dd2 (3.5 fold, 95% CI 3.0-4.1 fold), intermediate for NF54 (2.4 fold, 95% CI 2.0-3.0 fold) and HB3 (2.0 fold, 95% CI 1.6-2.5 fold), and least for 3D7 (1.5 fold, 95% CI 1.2-1.9 fold). All individual counts and assay values for all biological replicates are given in Supplementary Table S7.

The mean response of each parasite line to choline was estimated by the Mantel-Haenszel rate ratio (RR_M-H_), which analyses data across replicates under a simple fixed-effects model. With the exception of T9/96 which showed no response as noted above, the remaining four lines show overall increases in gametocyte conversion rates, but the mean responses varied among the different lines (Supplementary Table S7). The greatest mean response to choline was shown by line Dd2 (3.5 fold, 95% CI 3.0-4.1 fold), while 3D7 showed the lowest level of effect (1.5 fold, 95% CI 1.2-1.9 fold), and the other lines showed intermediate levels (for NF54 = 2.4 fold, 95% CI 2.0-3.0 fold; for HB3 = 2.0 fold, 95% CI 1.6-2.5 fold).

## Discussion

Analyses of multiple biological replicate assays are informative to characterise the range of sexual commitment rates in *P. falciparum*. A controlled comparison of two alternative methods showed that phenotyping was more sensitive and precise with an assay that measures proportions of sexually differentiating parasites by immunofluorescence staining of the Pfs16 marker expressed in early stage I gametocytes [15]. Although the sensitivity and precision of the methods differ, they give correlated measurements, and the lower proportions measured using a method that does not employ Pfs16 staining may be expected, given the requirement for parasites to successfully develop to stage II gametocytes over four days in culture before they are counted by morphological inspection of Giemsa-stained slides [33].

The recently culture-established Ghanaian clinical isolates analysed here had gametocyte conversion rates of between 3.3% and 12.2% per cycle, measured as means of results from at least six biological replicate assays for each isolate. These rates are higher than those previously estimated for clinical isolates in the first cycle *ex vivo* in Ghana with an assay which involves counting of stage II gametocytes, which indicated median rates less than 1% and only approximately a quarter of isolates having rates above 3% [33]. The lower *ex vivo* rates may be partly due to lower assay sensitivity compared to the Pfs16 staining method, as well as effects of plasma components such as variable lysophosphatidylcholine levels in some individuals inhibiting parasite sexual commitment [33]. It is notable that the isolates in the present study were cultured consistently in serum-free medium supplemented by Albumax-II for several months, demonstrating that use of serum in culture is not required to preserve the intrinsic capacity of parasites to show robust rates of sexual commitment over this period. Nevertheless, parasites with very high rates of sexual commitment might have been selected against, while eventual loss of sexual commitment in some cultures may occur over longer periods of culture.

Most of the diverse long-term laboratory-adapted lines had mean gametocyte conversion rates between 4.7% and 14.5% per cycle, and showed variation in conversion rate among different biological replicates. The only line that always had zero sexual conversion was F12 with a previously described non-functional *ap2-g* gene due to a nonsense mutation [6]. All other lines showed some detectable sexual conversion, although four lines had much lower rates (means of 0.3 to 1.6%) than the rest, likely due to loss-of-function mutations acquired in culture including deletion of the *gdv1* gene in lines T9/96 and D10 [36] which showed the lowest conversion rates. Loss of *gdv1* in two other laboratory lines was reportedly associated with no detectable gametocytes [12, 37], and it is possible that the low level of conversion in T9/96 and D10 reflects a small amount of commitment to gametocyte development post-invasion due to stochastic activation of *ap2-g*, independent of the main pathway involving *gdv1* [15].

The multiple replicate measurements of each *P. falciparum* line enable the identification of significant variation among recently derived clinical isolates from the same local endemic population, as well as among long-term laboratory-adapted lines that have diverse geographical origins. Given that modelling of data from experimental infections has indicated much lower rates of approximately 1% or less per cycle [20, 21], there may be a significant loss *in vivo* during the long period of development to circulating late stage gametocytes, or suppression of gametocyte conversion in these infections compared to culture.

Variation in gametocyte conversion rates among biological replicate assays was similar for recently established clinical isolates and most of the long-term laboratory adapted lines. Measured differences among replicates may reflect intrinsic parasite variation, stochastic or due to determinants yet unidentified, or could be due to assay conditions. Previous studies on a several different culture-adapted *P. falciparum* clones did not show as much variation between experimental replicates [6, 13, 28, 31], which may be due to tighter parasite synchronisation in those studies, whereas a broad window of synchronisation in the present study was intended to be similar to what may be sampled in peripheral blood (in which ring stage parasites may range in ages up to approximately 24 hours). The inter-replicate assay variation in this study might be partly due to the possibility that a proportion of parasites in some replicates were too young to stain with the Pf16 marker, although this was aimed to be minimised by inspecting parasite development to determine the exact time of assay completion. Although most parasites assayed would be mature enough for Pfs16 staining, some replicates may have parasites at least 36 hours post invasion when all stage I gametocytes would be Pfs16-positive [15, 17, 40], while other replicates may have parasites 24 hours post invasion when a majority but not all stage I gametocytes may already be Pfs16-positive [15]. Exact standardisation of parasite ages in culture of diverse isolates is not feasible, as isolates intrinsically vary in developmental cycle times [41, 42], but it is likely that tighter synchronisation of parasite cultures would reduce variation among replicates for any individual isolate. Assay precision may also be increased if detection assays are developed to enable antibody staining of sexually committed parasites before they express Pfs16, which may become possible as particular gene transcripts are expressed in committed ring stages [43], and at least one protein (GEXP02) is expressed earlier than Pfs16 [17].

Although a majority of the diverse laboratory-adapted parasite lines tested in this study retain an intact mechanism for gametocyte production, there were differences in responses to choline. The presence or absence of choline had no effect in any of eight replicate experiments on the T9/96 clone that has a culture-acquired deletion of *gdv1*, consistent with the requirement of this locus for mediating a response to induce enhanced levels of sexual commitment [12, 13]. In contrast, choline had a significant but variable effect on sexual commitment in each of the other parasite lines tested, each of which also showed substantial variation in responsiveness among biological replicates. Although a few individual replicate assays showed greater than 10 fold effects, mean suppressive effects of choline in each parasite line ranged from 1.5 fold (for 3D7) to 3.5 fold (for Dd2), more moderate than described in previous studies of parasite clones maintained in medium containing choline or serum before being switched to choline-free medium [13, 17, 31]. Gametocyte conversion rates of >30% have occasionally been reported elsewhere [28], which were not seen in the non-genetically engineered lines here, but only with overexpression of GDV1 in an engineered line. As all parasite lines including clinical isolates in this study were maintained in a standard medium with Albumax without choline supplementation, it is possible that higher gametocyte conversion rates were selected against, as they might not be sustained in continuous culture without using conditions to suppress conversion during long-term maintenance [31].

The results support the utility of choline in medium to suppress parasite sexual commitment under controlled environmental conditions (10, 20, 21), and notably show significant variation in the responsiveness of different parasite lines. It will be important to understand if there is intraspecific variation in natural regulation of GDV1 expression, particularly the control of gene-silencing antisense transcripts [13], and investigate the potential influence of naturally-occurring cis-variation at the sub-telomeric locus on chromosome 9 [33, 39, 44] distinct from the deletions disrupting function in some cultured lines [12, 36, 37, 45]. An important new insight into the effect of available choline has just been reported, indicating that sexual commitment is affected by competition between histone methyltransferases (involved in *ap2-g* gene silencing) and phosphoethanolamine methyltransferase (involved in *de novo* synthesis of phosphatidylcholine when choline availability is low) for the methyl donor S-adenosylmethionine, which could have implications for future studies on inter-isolate variation in rates of sexual commitment [32]. Investigation of parasites in natural populations requires assay precision and experimental replication to be considered where possible alongside statistical power from increased numbers of clinical samples, whether for candidate gene investigations or larger scale scanning for associations with unlinked genomic variation in parasite populations [46].

## Materials and Methods

### Ethics Approval

Approval for sampling of clinical isolates for culture experimental analysis was granted by the Ethics committees of the Ghana Health Service, the Noguchi Memorial Institute for Medical Research at the University of Ghana, the Navrongo Health Research Centre and the London School of Hygiene and Tropical Medicine, as previously described for the collection [35] from which specific isolates were selected in this analysis.

### Plasmodium falciparum laboratory-adapted clones and clinical isolates

The genetically modified *P. falciparum* cloned line 3D7/iGP_D9 was used to compare the sensitivity and precision of gametocyte conversion rate assays, as artificially elevated sexual commitment is selectively inducible in this line by overexpression of GDV1 fused to a destabilization domain (DD) which maintains activity in the presence of Shield-1 reagent [34].

Thirteen different *P. falciparum* long-term laboratory-adapted lines that have not undergone targeted genetic modification were studied. Each of these was either previously derived by cloning or is a line that has an apparently monoclonal single-genome sequence, and a putative origin based on the geographical location of the original isolation: NF54 (a strain of ‘airport malaria’ from The Netherlands with genome sequence similarity to African isolates), 3D7 (the *P. falciparum* reference genome strain originally cloned from NF54 and maintained in culture for many years) [47], F12 (derived by cloning from cultured 3D7 and having a nonsense mutant *ap2-g* gene) [6], D10 (Papua New Guinea), D6 (Sierra Leone), Dd2 (Southeast Asia), FCC2 (China), HB3 (Honduras), R033 (Ghana), Palo Alto (Uganda), 7G8 (Brazil), GB4 (Ghana) and T9/96 (Thailand). All lines were thawed from pre-existing frozen stocks with distinct identities verified by previous targeted genotyping and sequencing [39, 48] and were not re-cloned prior to this study, and each line was cultured for at least one week after thawing prior to any assay, all culture being conducted using a commercially-prepared serum-free medium supplemented with lipid-rich bovine serum albumen (Gibco RPMI with 0.5% AlbuMAX™ II lot number 2177881). This consistent medium preparation had very minimal choline (less than 0.02 mM choline chloride) which would not suppress parasite sexual commitment, unlike high concentrations of choline (2 mM as used for some assays described in a separate section below) or serum which has variable components between batches.

Six *P. falciparum* clinical isolates from Ghana with single-genome unmixed infections were selected from a larger panel of infections sampled from children with clinical malaria in Navrongo (in the Upper East Region of the country), in which parasite asexual multiplication rate variation was previously studied [35]. These isolates were cultured with the same batch of commercially-prepared serum-free medium with 0.5% AlbuMAX™ II as for the laboratory-adapted parasite lines, ensuring the same batch was used throughout all experiments so that no biological variation should be due to media components, and these clinical isolates were not cultured with added serum at any time as they had been cultured with 0.5% AlbuMAX™ II since isolation [35]. Clinical isolates with mixed genome-infections were purposefully not selected for inclusion in this study, as their sexual commitment rates would be complex to interpret due to potential changes in mixed genomic composition over time [35].

### Quantification of sexual commitment rates by measuring gametocyte conversion

Multiple biological replicate preparations of *P. falciparum* long-term laboratory-adapted lines and more recently culture-established clinical isolates were each cultured in human erythrocytes at 3% hematocrit in RPMI 1640 media supplemented with AlbuMAX™ II and incubated at 37°C in atmospheric air with 5% CO_2_. Prior to assay, each asynchronous parasite culture was synchronized to within a 24-hour window of intra-erythrocytic asexual stage development, intended to correspond approximately to the parasite age range that would be typically sampled in peripheral blood of patients, using the following two-step technique. Parasites containing haemozoin, predominantly schizont and late trophozoite stages, were first positively selected on MACS® LD magnetic columns [49], and after culture for 24 hours a second MACS purification step was performed from which the flow through erythrocytes (containing predominantly ring stage parasites) were collected and put back into culture, which then contained exclusively ring stage parasites as any gametocytes or residual mature asexual parasites were removed on the column. This broad synchronization scheme is illustrated in Supplementary Figure S1. For the 3D7/iGP parasite line the effect of GDV1-DD overexpression was tested by adding Shield-1 reagent to this preparation of ring stages (1 µM compared to none, with intermediate concentration treatments of 0.05, 0.15 and 0.5 µM Shield-1 also being tested in some assays), with Shield-1 being retained in the culture throughout the cycle.

Two methods were used for measuring the gametocyte conversion rate:

#### Method 1. Counting of the proportion of stage I gametocytes compared to all parasites developing in the first cycle post-invasion using anti-Pfs16 staining

This method involves distinguishing very early gametocytes from asexual parasites by detection of the gametocyte marker Pfs16 protein [15, 40, 50]. After two rounds of MACS column purification to obtain ring stages followed by a further 24 hours of growth, when a culture with a broad synchronicity containing schizonts was obtained, parasites were allowed to invade erythrocytes and were cultured for a period of between 38 and 46 hours, the exact time of harvesting of each culture being decided on the basis of microscopical examination of Giemsa stained slides indicating that most post-invasion parasites were clearly beyond the ring stage of development and therefore should be either trophozoites or stage I gametocytes. The harvested post-invasion parasites were examined by fluorescence microscopy by Pfs16 staining of early gametocytes and DAPI staining of all parasites. Stage I gametocytes were counted as those staining positive for Pfs16 and showing typical round morphology similar to trophozoites (whereas any Pfs16-positive parasites with differentiated morphology of stage II gametocytes potentially residual from induction prior to the previous cycle were not counted). The assay enables most sexually-developing parasites to be counted in comparison to the total of post-invasion parasites, on the basis that a majority of early stage I gametocytes express Pfs16 after 24 hours post invasion [15] as a proportion of all gametocytes which eventually express [40], with only a minority of sexual parasites likely not to be counted due to being too young. Parasite cultures harvested for this purpose were washed in phosphate buffered saline (PBS) with 3% bovine serum albumen (BSA), resuspended to 2% hematocrit, spotted onto multiwell slides (Hendley, Essex, UK), air-dried and stored at -80°C. Prior to immunofluorescence staining, slides were fixed with 4% paraformaldehyde in PBS for 30 min and permeabilized with 0.1% Triton X-100 in PBS for 10 min. The slides were incubated for 30 minutes at room temperature with a monoclonal α-Pfs16 murine antibody 93A3A2 [50], diluted 1:2000 in PBS with 3% BSA. Alexa Fluor® 594-conjugated anti-mouse IgG (Invitrogen, A11032) was used as secondary antibody, diluted 1:1000 in PBS with 3% BSA and incubated for 30 minutes at room temperature. Slides were mounted with coverslips using DAPI-containing Prolong™ Diamond antifade mountant (ThermoFisher Scientific), and parasite counting was performed using a Zeiss CCD fluorescence microscope, with identical settings used for each experiment. An average of approximately 1000 parasites were counted to determine proportions expressing Pfs16 in each measurement, and where the density of parasites did not allow this a quality control was performed to exclude any assay data that did not have a minimum of at least 300 parasites counted.

#### Method 2.Counting of developing gametocytemia compared to all parasites at the ring stage culture

This method estimates the proportion of parasites that produce gametocytes developing to morphological stage II, by comparing with the ring parasitaemia counted 4 days previously at the beginning of the parasite developmental cycle. The technique is based on a protocol that uses Giemsa staining and does not require immunofluorescence staining of an early gametocyte marker [33]. To initiate this assay, after two MACS column purifications and when a broadly synchronous culture containing a majority of schizonts were obtained, parasites were allowed to re-invade fresh erythrocytes and cultured until ring stage development was observed. Then, ring stage parasitaemia was assessed by visual examination of thin blood smears of each culture to determine the proportion of erythrocytes containing parasites (referred to as D0 parasitaemia), ensuring that at least 1,000 erythrocytes were counted. The cultures were then treated with 50 mM N-acetyl-D-glucosamine (GlcNAc) to prevent further development of asexual parasites [51]. After 4 days, the stage II gametocytemia was assessed by counting of Giemsa-stained thin blood smears (referred to as D4 gametocytemia), ensuring that at least 10,000 erythrocytes were counted. The gametocyte conversion rate was calculated by dividing D4 stage II gametocytemia (as a proportion of erythrocytes infected) by D0 ring parasitaemia (as a proportion of erythrocytes infected), yielding a ‘ratio-of-ratio’ proportion with 95% confidence intervals.

For both methods, all assays were set up at 3% haematocrit in a culture volume of 5 ml (static in 6-well culture plates), with parasitaemia levels at the start of each replicate assay being between 0.5 -2.5% erythrocytes infected, to allow sufficient numbers of parasites for high precision counting while minimising stress on parasites that may be induced if cultures have higher parasitaemia.

### Quantification of gametocyte conversion rate using culture media without choline

Parasites were prepared for assays in the same manner as described in the previous section, except that they were split after the second MACS column purification (at ring stage in the cycle before gametocyte counting) into separate wells with culture media (RPMI 1640 with 25 mM HEPES, 100 µM Hypoxanthine, 1 mM L-Glutamine, 0.39% fatty acid-free BSA, 30 mM Palmitic acid, 30 mM Oleic acid) either supplemented with 2 mM choline or lacking choline, based on a previously described protocol [31]. Parasites were allowed to grow until schizont stage and to invade fresh erythrocytes, without changing the culture medium. After 38-46 hours post invasion, parasites were collected and fixed for analysis of gametocyte conversion by differential counting using Pfs16 staining by immunofluorescence microscopy as well as DAPI staining as described above for Method 1, with an average of approximately 1000 parasites being counted for each individual measurement. Between six and eight replicate assays were performed for each parasite line, the effect within each replicate calculated as the rate ratio (with 95% confidence intervals) of the gametocyte conversion rate in choline-free medium compared with the conversion rate in medium containing 2 mM choline.

### Statistical analyses

Spearman’s ρ (rho) coefficient was used to evaluate the strength and significance of the rank correlation between results from the two different gametocyte conversion rate assays performed in parallel. Statistical comparisons of gametocyte conversion rate variation between different *P. falciparum* lines were performed using the Mann-Whitney test on the multiple replicate assay measurements of each line. Each different parasite line was assayed with a mean of more than six biological replicate cultures, most individual lines having at least six and some up to nine biological replicates performed. Each individual biological replicate data point had 95% confidence intervals calculated for the proportions of parasites showing sexual commitment on the basis of the raw numerical counts performed, enabling accurate estimation of within-assay sampling variation, and all count data are provided for verification and to enable any future meta-analyses. For each parasite line, the mean effect of modified culture media conditions was estimated by calculating the Mantel-Haenszel rate ratio (RR_M-H_) across biological replicate assays, based on a simple fixed-effects model. Statistical analyses were performed using Prism version 9 and Epi-Info version 7 software.

## Acknowledgements

This study was funded by Project Grant MR/S009760/1 from the UK Medical Research Council to DJC and DAB. Original sampling of clinical isolates was supported by a Royal Society Africa Award AA110050 to GAA and DJC. TSV received funding from the Swiss National Science Foundation grant BSCGI0_157729. We are grateful to all colleagues who supported the research environment, including sharing of laboratory facilities and equipment, enabling continuous parasite culture and experimentation as well as storage and shipment of samples and reagents.

