## Supplementary for "Variation in *Plasmodium falciparum* sexual commitment rates and responses to environmental modification"

### Supplementary Information

Lindsay B. Stewart [1a], Aline Freville [1a], Till S. Voss [2], David A. Baker [1], Gordon A. Awandare [3], David J. Conway [1*]

[1] Department of Infection Biology, Faculty of Infectious and Tropical Diseases, London School of Hygiene and Tropical Medicine, London WC1E 7HT, UK

[2] Department of Medical Parasitology and Infection Biology, Swiss Tropical and Public Health Institute, and University of Basel, Switzerland

[3] West African Centre for Cell Biology of Infectious Pathogens (WACCBIP), Department of Biochemistry, Cell and Molecular Biology, University of Ghana, Accra, Ghana

[a] These authors contributed equally to the research

**Supplementary Table S1.** Gametocyte conversion rate (GCR) estimates (with 95% confidence intervals) and all numerical count data for two different methods with multiple experimental replicates applied to *P. falciparum* line 3D7/iGP under elevated GDV1 expression with Shield-1 reagent (+SHLD) and control (-SHLD) conditions.

| | | Method 1 | | | Method 2 | | |
|---|---|---|---|---|---|---|---|
| **Replicate** | **Condition** | **Pfs16 +ve** | **DAPI +ve** | **Pfs16 GCR (95%CI)** | **D0 % Parasit.** | **D4 % Parasit.** | **D0D4 GCR (95% CI)** |
| 1 | -SHLD | 84 | 921 | 9.12 (7.4 - 11.2) | 10.1 (101/999) | 0.3 (26/8262) | 3.4 (2.2 -5.2) |
| | +SHLD | 474 | 826 | 57.4 (54.0 - 60.7) | 9.8 (98/999) | 1.9 (117/6124) | 20.9 (16.0 - 27.3) |
| 2 | -SHLD | 104 | 1171 | 8.9 (7.4 - 10.7) | 7.2 (67/932) | 0.2 (15/8210) | 2.7 (1.6 - 4.7) |
| | +SHLD | 262 | 978 | 26.8 (24.1 - 29.7) | 3.0 (29/970) | 0.4 (49/13415) | 12.5 (7.9 - 19.8) |
| 3 | -SHLD | 41 | 926 | 4.4 (3.3 - 6.0) | 5.6 (53/946) | 0.1 (8/9305) | 1.6 (2.8 - 3.4) |
| | +SHLD | 150 | 595 | 25.2 (21.9 - 28.9) | 4.4 (42/957) | 0.4 (36/8625) | 9.9 ( 6.3 - 15.4) |
| 4 | -SHLD | 47 | 1043 | 4.5 (3.4 - 6.0) | 5.0 (48/951) | 0.4 (30/7278) | 8.5 (5.4 - 13.5) |
| | +SHLD | 267 | 701 | 38.1 (34.6 - 41.7) | 4.4 (42/952) | 1.0 (77/8070) | 20.1 (14.0 - 28.8) |

**Supplementary Table S2.** Estimates of gametocyte conversion rate (GCR) for 17 assays of the *P. falciparum* inducible 3D7/iGP_D9 line in which two different methods were performed in parallel for comparison (as detailed in Materials and Methods). Varying concentrations of Shield-1 (0.055 – 1.0 µM, and none) were used for the assays. Data are plotted and analysed in Figure 2, showing an overall correlation between the methods, with Method 1 being generally more sensitive (and more precise with narrower confidence intervals as shown here).

| Independent assays of *P. falciparum* line | Method 1 Pfs16 GCR (95%CI) | Method 2 D0/D4 GCR (95%CI) |
|---|---|---|
| iGP - SHLD | 9.1 (7.4 - 11.2) | 3.4 (2.2 - 5.3) |
| iGP +SHLD (1.0) | 57.4 (54.0 - 60.7) | 20.9 (16.0 - 27.31) |
| iGP - SHLD | 8.9 (7.4 - 10.7) | 2.7 (1.6 - 4.8) |
| iGP + SHLD (0.055) | 14.2 (12.3 - 16.4) | 4.0 (2.2 - 7.2) |
| iGP + SHLD (0.15) | 16.8 (14.8 - 19.1) | 2.4 (1.2 (4.9) |
| iGP + SHLD (0.5) | 17.4 (15.3 - 19.7) | 2.6 (1.3 - 5.2) |
| iGP +SHLD (1.0) | 26.8 (24.1 - 30.0) | 12.5 (8.0 - 19.8) |
| iGP - SHLD | 4.4 (3.4 - 7.0) | 1.6 (0.8 - 3.4) |
| iGP + SHLD (0.055) | 20.5 (17.9 - 23.4) | 7.2 (4.6 - 11.3) |
| iGP + SHLD (0.15) | 16.1 (13.8 - 18.7) | 4.7 (3.2 - 6.9) |
| iGP + SHLD (0.5) | 26.5 (23.5 - 29.6) | 5.1 (3.2 - 8.2) |
| iGP +SHLD (1.0) | 25.2 (21.9 - 28.9) | 9.9 (6.4 - 15.4) |
| iGP - SHLD | 4.5 (3.4 - 6.9) | 8.5 (5.4 - 13.5) |
| iGP + SHLD (0.055) | 8.9 (7.2 - 11.2) | 10.7 (7.3 - 15.7) |
| iGP + SHLD (0.15) | 22.3 (19.2 - 25.8) | 8.9 (6.1 - 12.8) |
| iGP + SHLD (0.5) | 37.4 (34.0 - 41.0) | 17.5 (12.9 - 23.8) |
| iGP +SHLD (1.0) | 38.1 (34.6 - 41.7) | 20.1 (14.0 - 28.8) |

**Supplementary Table S3.** Multiple biological replicate assays of gametocyte conversion rates (GCR) for each of 6 cultured *P. falciparum* clinical isolates from Navrongo, Ghana.

| Isolate | Replicate | RBC Batch | Days in culture | % Parasit. | Pfs16 +ve | DAPI +ve | GCR (95%CI) |
|---|---|---|---|---|---|---|---|
| 289 | 1 | 12 | 92 | 1.4 | 38 | 961 | 4.0 (2.9 - 5.4) |
| | 2 | 12 | 99 | 0.7 | 9 | 870 | 1.0 ( 0.6 - 2.0) |
| | 3 | 12 | 99 | 0.9 | 23 | 740 | 3.1 (2.1 - 4.6) |
| | 4 | 13 | 106 | 0.9 | 53 | 946 | 5.6 (4.3 - 7.3) |
| | 5 | 13 | 106 | 0.8 | 28 | 971 | 2.9 (2.0 - 4.1) |
| | 6 | 15 | 125 | 0.8 | 28 | 939 | 3.0 (2.1 - 4.3) |
| | *Mean (SD)* | | | | | | *3.3 (SD = 1.5)* |
| | 1 | 12 | 83 | 1.0 | 48 | 951 | 5.0 (3.8 - 6.6) |
| | 2 | 12 | 83 | 0.8 | 45 | 954 | 4.7 (3.5 - 6.3) |
| | 3 | 13 | 91 | 1.2 | 34 | 965 | 3.5 (2.5 - 4.9) |
| | 4 | 13 | 91 | 0.6 | 26 | 973 | 2.7 (1.8 - 3.9) |
| | 5 | 15 | 110 | 0.6 | 69 | 999 | 6.9 (5.5 - 8.7) |
| 272 | 6 | 15 | 110 | 0.6 | 51 | 880 | 5.8 (4.4 - 7.5) |
| | *Mean (SD)* | | | | | | *4.8 (SD = 1.5)* |
| | 1 | 1 | 67 | 0.8 | 66 | 677 | 9.7 (7.7 - 12.2) |
| | 2 | 12 | 62 | 0.9 | 34 | 965 | 3.5 (2.5 - 4.9) |
| | 3 | 12 | 62 | 0.7 | 58 | 882 | 6.6 (5.1 - 8.4) |
| 292 | 4 | 13 | 69 | 2.3 | 63 | 936 | 6.7 (5.3 - 8.5) |
| | 5 | 13 | 69 | 1.1 | 46 | 953 | 4.8 (3.6 - 6.4) |
| | 6 | 15 | 88 | 0.5 | 49 | 939 | 5.2 (4.0 - 6.8) |
| | 7 | 15 | 88 | 0.8 | 19 | 564 | 3.4 (2.2 - 5.2) |
| | *Mean (SD)* | | | | | | *5.7 (SD = 2.2)* |
| | 1 | 1 | 67 | 1.7 | 109 | 1841 | 5.9 (4.9 - 7.1) |
| | 2 | 12 | 55 | 0.8 | 120 | 878 | 13.7 (11.6 - 16.1) |
| 293 | 3 | 13 | 69 | 0.9 | 118 | 881 | 13.4 (11.3 - 15.8) |
| | 4 | 13 | 69 | 1.3 | 118 | 883 | 13.4 (11.3 - 15.8) |
| | 5 | 13 | 76 | 1.0 | 39 | 962 | 4.1 (3.0 - 5.5) |
| | 6 | 13 | 76 | 1.0 | 41 | 617 | 6.6 (4.9 - 8.9) |
| | *Mean (SD)* | | | | | | *9.5 (SD = 4.4)* |
| | 1 | 12 | 48 | 1.5 | 116 | 883 | 13.1 (11.1 - 15.5) |
| | 2 | 12 | 55 | 1.0 | 31 | 968 | 3.2 (2.3 - 4.5) |
| | 3 | 12 | 55 | 0.6 | 65 | 632 | 10.3 (8.2 - 12.9) |
| 296 | 4 | 13 | 62 | 1.2 | 94 | 905 | 10.4 (8.6 - 12.5) |
| | 5 | 13 | 62 | 1.1 | 84 | 915 | 9.2 (7.5 - 11.2) |
| | 6 | 13 | 69 | 1.0 | 93 | 906 | 10.3 (8.5 - 12.4) |
| | 7 | 13 | 69 | 1.0 | 118 | 887 | 13.3 (11.2 - 15.7) |
| | *Mean (SD)* | | | | | | *10.0 (SD = 3.4)* |
| | 1 | 1 | 55 | 1.0 | 156 | 1406 | 11.1 (9.6 - 12.9) |
| | 2 | 2 | 64 | 2.2 | 103 | 360 | 28.6 (24.2 - 33.5) |
| | 3 | 2 | 64 | 1.5 | 209 | 794 | 26.3 (23.5 - 29.6) |
| | 4 | 3 | 71 | 0.7 | 131 | 907 | 14.4 (12.3 - 16.9) |
| 280 | 5 | 3 | 71 | 1.0 | 109 | 787 | 13.9 (11.6 - 16.4) |
| | 6 | 3 | 78 | 1.4 | 27 | 973 | 2.8 (1.9 - 4.0) |
| | 7 | 3 | 78 | 0.6 | 49 | 884 | 5.5 (4.2 - 7.3) |
| | 8 | 4 | 85 | 1.4 | 12 | 998 | 1.2 (0.7 - 2.1) |
| | 9 | 4 | 85 | 0.6 | 56 | 943 | 5.9 (4.6 - 7.6) |
| | *Mean (SD)* | | | | | | *12.2 (SD = 9.8)* |

The table shows exact counts and 95% confidence intervals of the GCR estimates using Method 1 (with Pfs16 staining) as plotted and analysed in Figure 3. The number of days in culture is shown as well as the % parasitaemia measured in the cycle before. The assay replicates are presented in the order in which they were performed (the duration of time in culture occasionally does not exactly follow the same sequential temporal order for Isolates 292 and 293 as an intermediate cryopreservation and thawing of the isolates was performed). Erythrocytes from 15 different erythrocyte donors used at various times throughout the series of experiments shown here and in Supplementary Table S5 are numbered (RBC Batch), and inspection of the data indicates that these did not determine the variation between assay replicates.

**Supplementary Table S4.** Pairwise tests for significant differences in gametocyte conversion rates between different Ghanaian *P. falciparum* clinical isolates.

| Clinical Isolates | 272 | 280 | 289 | 292 | 293 | 296 |
|---|---|---|---|---|---|---|
| 272 | | | | | | |
| 280 | 0.1447 | | | | | |
| 289 | 0.1797 | 0.0663 | | | | |
| 292 | 0.5594 | 0.2523 | 0.0350* | | | |
| 293 | 0.0649 | 0.7756 | 0.0043** | 0.1375 | | |
| 296 | 0.0221* | 0.7577 | 0.0047** | 0.0379* | 0.7308 | |

P values are from Mann-Whitney comparisons of all of the individual biological replicate measures for each of the six isolates (as shown in Supplementary Table S3). Statistically significant comparisons are highlighted in green.

**Supplementary Table S5.** Multiple biological replicate assays of gametocyte conversion rates (GCR) for each of 13 long-term culture adapted *P. falciparum* lines of diverse origins.

| Line | Replicate | RBC batch | Pfs16 +ve | DAPI +ve | GCR (95% CI) |
|---|---|---|---|---|---|
| F12 | 1 | 3 | 0 | 1000 | 0 (0) |
| | 2 | 3 | 0 | 1000 | 0 (0) |
| | 3 | 4 | 0 | 1023 | 0 (0) |
| | 4 | 10 | 0 | 394 | 0 (0) |
| | 5 | 11 | 0 | 600 | 0 (0) |
| | 6 | 11 | 0 | 711 | 0 (0) |
| | 7 | 11 | 0 | 380 | 0 (0) |
| | *Mean (SD)* | | | | 0 (SD = 0) |
| D10 | 1 | 2 | 0 | 515 | 0 (0 ) |
| | 2 | 2 | 1 | 999 | 0.1 ( 0 - 5.6) |
| | 3 | 2 | 10 | 1177 | 0.8 (0.5 - 1.6) |
| | 4 | 3 | 2 | 997 | 0.2 (0.1 - 0.7) |
| | 5 | 3 | 2 | 1000 | 0.2 (0.1 -0.7) |
| | 6 | 9 | 3 | 997 | 0.3 (0.1 - 0.9) |
| | 7 | 9 | 2 | 997 | 0.2 ( 01 - 0.7) |
| | *Mean (SD)* | | | | 0.26 (SD = 0.27) |
| T996 | 1 | 12 | 6 | 993 | 0.6 (0.3 - 1.3) |
| | 2 | 13 | 2 | 733 | 0.3 (0.1 - 1.0) |
| | 3 | 13 | 3 | 992 | 0.3 (0.1 - 0.9) |
| | 4 | 13 | 2 | 703 | 0.3 (0.1 - 1.0) |
| | 5 | 14 | 2 | 997 | 0.2 (0 - 0.7) |
| | 6 | 14 | 3 | 433 | 0.7 (0.2 - 2.0) |
| | *Mean (SD)* | | | | 0.39 (SD = 0.20) |
| Palo Alto | 1 | 6 | 12 | 987 | 1.2 (0.7 - 1.2) |
| | 2 | 7 | 14 | 514 | 2.7 (1.6 - 4.5) |
| | 3 | 7 | 5 | 321 | 1.6 (0.7 - 3.6) |
| | 4 | 9 | 1 | 998 | 0.1 (0 - 0.6) |
| | 5 | 9 | 3 | 996 | 0.3 (0.1 - 0.9) |
| | 6 | 9 | 1 | 998 | 0.1 (0 - 0.6) |
| | *Mean (SD)* | | | | 1.00 (SD = 1.04) |
| 3D7 | 1 | 2 | 20 | 980 | 2.0 (1.3 - 3.1) |
| | 2 | 3 | 3 | 994 | 0.3 (0.1 - 0.9) |
| | 3 | 3 | 12 | 979 | 1.2 (0.7 -2.1) |
| | 4 | 4 | 20 | 977 | 2.0 (1.3 -3.1) |
| | 5 | 4 | 17 | 916 | 1.9 (1.2 - 3.0) |
| | 6 | 12 | 20 | 885 | 2.3 (1.5 - 3.5) |
| | 7 | 12 | 16 | 983 | 1.6 (1.0 - 2.6) |
| | *Mean (SD)* | | | | 1.62 (SD = 0.67) |

| Line | Replicate | RBC batch | Pfs16 +ve | DAPI +ve | GCR (95% CI) |
|---|---|---|---|---|---|
| D6 | 1 | 5 | 83 | 920 | 9.0 (7.3 -11.1) |
| | 2 | 5 | 16 | 983 | 1.6 (1.0 - 2.6) |
| | 3 | 5 | 6 | 885 | 0.7 (0.3 - 1.5) |
| | 4 | 9 | 28 | 559 | 5.0 (3.5 - 7.1) |
| | 5 | 9 | 86 | 905 | 9.5 (7.8 - 11.6) |
| | 6 | 9 | 26 | 973 | 2.7 (1.8 - 3.9) |
| | 7 | 9 | 50 | 950 | 5.3 (4.0 - 6.9) |
| | *Mean (SD)* | | | | 4.70 (SD = 3.36) |
| GB4 | 1 | 6 | 22 | 978 | 2.2 (1.5 - 3.4) |
| | 2 | 6 | 85 | 914 | 9.3 (7.6 -11.4) |
| | 3 | 6 | 92 | 917 | 10.0 (8.3 - 12.2) |
| | 4 | 6 | 37 | 964 | 3.8 (2.8 - 5.3) |
| | 5 | 7 | 59 | 940 | 6.3 (4.9 - 8.0) |
| | 6 | 7 | 16 | 983 | 1.6 (1.0 - 2.6) |
| | 7 | 9 | 34 | 969 | 3.5 (2.5 - 4.9) |
| | *Mean (SD)* | | | | 5.26 (3.35) |
| Dd2 | 1 | 2 | 130 | 902 | 14.4 (12.3 - 16.9) |
| | 2 | 3 | 4 | 995 | 0.4 (0.2 - 1.0) |
| | 3 | 3 | 9 | 986 | 0.9 (0.5 - 1.7) |
| | 4 | 4 | 46 | 953 | 4.8 (3.6 - 6.4) |
| | 5 | 4 | 84 | 916 | 9.2 (7.5 - 11.2) |
| | 6 | 12 | 23 | 532 | 4.3 (2.9 -6.4) |
| | 7 | 12 | 35 | 885 | 4.0 (2.9 - 5.5) |
| | *Mean (SD)* | | | | 5.42 (SD = 4.89) |
| HB3 | 1 | 2 | 50 | 691 | 7.2 (5.5 - 9.4) |
| | 2 | 2 | 103 | 1000 | 10.3 (8.6 - 12.3) |
| | 3 | 9 | 57 | 942 | 6.1 (4.7 -7.8) |
| | 4 | 9 | 56 | 944 | 5.9 (4.6 -7.6) |
| | *Mean (SD)* | | | | 6.87 (SD = 2.32) |
| NF54 | 1 | 2 | 225 | 1022 | 22.0 (19.6 - 24.7) |
| | 2 | 2 | 182 | 1013 | 18.0 (15.7 - 20.5) |
| | 3 | 3 | 73 | 926 | 7.9 (6.3 - 9.8) |
| | 4 | 3 | 54 | 925 | 5.8 (4.5 - 7.5) |
| | 5 | 13 | 41 | 619 | 6.6 (4.9 - 8.9) |
| | 6 | 13 | 11 | 975 | 1.1 (0.6 - 2.0) |
| | 7 | 13 | 27 | 975 | 2.8 (1.9 - 4.0) |
| | *Mean (SD)* | | | | 9.18 (SD = 7.83) |

| Line | Replicate | RBC Batch | Pfs16 +ve | DAPI +ve | GCR (95% CI) |
|---|---|---|---|---|---|
| RO33 | 1 | 5 | 153 | 861 | 17.8 (15.4 - 20.5) |
| | 2 | 5 | 68 | 943 | 7.2 (5.7 - 9.0) |
| | 3 | 7 | 70 | 932 | 7.5 (6.0 - 9.4) |
| | 4 | 7 | 92 | 910 | 10.1 (8.3 - 12.2) |
| | 5 | 10 | 72 | 921 | 7.8 (6.3 - 9.7) |
| | 6 | 10 | 124 | 875 | 14.2 (12.0 - 16.6) |
| | *Mean (SD)* | | | | 10.77 (SD = 4.31) |
| FCC2 | 1 | 5 | 211 | 785 | 26.9 (23.9 - 30.1) |
| | 4 | 5 | 147 | 835 | 17.6 (15.2 - 20.3) |
| | 5 | 5 | 16 | 484 | 3.3 (2.0 - 5.3) |
| | 6 | 5 | 23 | 503 | 4.6 (3.1 - 6.8) |
| | 7 | 10 | 69 | 935 | 7.4 (5.9 - 9.2) |
| | 8 | 10 | 47 | 952 | 4.9 (3.7 - 6.5) |
| | *Mean (SD)* | | | | 10.78 (SD = 9.44) |
| 7G8 | 1 | 6 | 159 | 856 | 18.6 (16.1 - 21.3) |
| | 2 | 6 | 199 | 803 | 24.8 (21.9 - 27.9) |
| | 3 | 10 | 135 | 452 | 29.9 (25.8 - 34.2) |
| | 4 | 10 | 24 | 339 | 7.1 (4.8 - 10.3) |
| | 5 | 10 | 9 | 456 | 2.0 (1.0 - 3.7) |
| | 6 | 11 | 46 | 953 | 4.8 (3.6 - 6.4) |
| | *Mean (SD)* | | | | 14.52 (SD = 11.52) |

Exact counts and 95% confidence intervals of the GCR estimates were performed using Method 1 (with Pfs16 staining) as plotted and analysed in Figure 4. Erythrocytes from fifteen different erythrocyte donors used at various times throughout the series of experiments shown here and in Supplementary Table S3 are numbered (RBC Batch), and inspection of the data shows that these did not determine the variation between assay replicates.

**Supplementary Table S6.** Pairwise tests for significant differences in gametocyte conversion rates between different *P. falciparum* laboratory lines. P values are from Mann-Whitney comparisons of all of the individual biological replicate measures for each of the 13 lines (as shown in Supplementary Table S5). Statistically significant comparisons are highlighted in green.

| Lab Isolates | F12 | D10 | T9/96 | Palo Alto | 3D7 | D6 | GB4 | Dd2 | HB3 | NF54 | RO33 | 7G8 | FCC2 |
|---|---|---|---|---|---|---|---|---|---|---|---|---|---|
| F12 | | | | | | | | | | | | | |
| D10 | 0.0047** | | | | | | | | | | | | |
| T9/96 | 0.0022** | 0.1696 | | | | | | | | | | | |
| Palo Alto | 0.0022** | 0.2459 | 0.6082 | | | | | | | | | | |
| 3D7 | 0.0012** | 0.0017** | 0.0052** | 0.1521 | | | | | | | | | |
| D6 | 0.0012** | 0.0012** | 0.0023** | 0.0216* | 0.0781 | | | | | | | | |
| GB4 | 0.0012** | 0.0006*** | 0.0012** | 0.0047** | 0.0117* | 0.6457 | | | | | | | |
| Dd2 | 0.0012** | 0.0012** | 0.0047** | 0.0338* | 0.1282 | >0.9999 | >0.9999 | | | | | | |
| HB3 | 0.0048** | 0.0030** | 0.0095** | 0.0061** | 0.0012** | 0.2303 | 0.4121 | 0.3124 | | | | | |
| NF54 | 0.0012** | 0.0006*** | 0.0012** | 0.0082** | 0.0175* | 0.3176 | 0.535 | 0.3176 | 0.9273 | | | | |
| RO33 | 0.0022** | 0.0012** | 0.0022** | 0.0022** | 0.0012** | 0.035* | 0.035* | 0.0734 | 0.026* | 0.4452 | | | |
| 7G8 | 0.0022** | 0.0012** | 0.0022** | 0.0043** | 0.0082** | 0.1807 | 0.1807 | 0.1101 | 0.3939 | 0.4452 | >0.9999 | | |
| FCC2 | 0.0022** | 0.0012** | 0.0022** | 0.0022** | 0.0012** | 0.336 | 0.2949 | 0.2343 | 0.9143 | 0.9452 | 0.3939 | 0.6991 | |

**Supplementary Table S7.** Numerical counts and calculated values of assays measuring the effect of choline-free medium on gametocyte conversion rates (GCR) of five different *P. falciparum* lines. The effect of each biological replicate assay is shown as a Rate Ratio with 95% confidence intervals, based on the GCR with and without choline. Mantel-Haenszel adjusted mean Rate Ratios show the overall effects across all replicates with 95% confidence intervals.

| Biological Replicate Assays | Condition | Pfs16 +ve | Pfs16 -ve | Rate Ratio (± 95%CI) | GCR (± 95%CI) | Adjusted mean Rate Ratio (± 95%CI) |
|---|---|---|---|---|---|---|
| | | | ***P. falciparum* line HB3** | | | |
| R1 | -choline | 16 | 967 | 2.03 (0.93 - 5.46) | 1.6 ( 1.0 - 2.6) | **1.98 (1.55 - 2.52)** |
| | + choline | 7 | 864 | | 0.8 (0.4 - 1.6) | |
| R2 | -choline | 29 | 943 | 3.69 (1.69 - 8.04) | 3.0 (2.1 - 4.3) | |
| | + choline | 8 | 983 | | 0.8 (0.4 - 1.6) | |
| R3 | -choline | 34 | 902 | 12.05 (3.71 - 39.13) | 3.6 (2.6 -5.0) | |
| | + choline | 3 | 993 | | 0.3 (0.1 - 0.9) | |
| R4 | -choline | 36 | 929 | 2.01 (1.15 - 3.51) | 3.7 (2.7 - 5.1) | |
| | + choline | 18 | 950 | | 1.9 (1.2 - 2.9) | |
| R5 | -choline | 24 | 954 | 1.26 (0.70 - 2.30) | 2.5 (1.7 - 3.6) | |
| | + choline | 19 | 961 | | 1.9 (1.2 - 3.0) | |
| R6 | -choline | 14 | 970 | 1.00 (0.48 - 2.09) | 1.4 (0.8 - 2.4) | |
| | + choline | 14 | 971 | | 1.4 (0.8 - 2.4) | |
| R7 | -choline | 20 | 959 | 0.95 (0.52 - 1.74) | 2.0 (1.3 - 3.1) | |
| | + choline | 21 | 957 | | 2.1 (1.4 - 3.3) | |
| R8 | -choline | 19 | 962 | 2.40 1.06 - 5.45) | 1.9 (1.2-3.0) | |
| | + choline | 8 | 983 | | 0.8 (0.4 - 1.6) | |
| | | | ***P. falciparum* line Dd2** | | | |
| R1 | -choline | 194 | 619 | 5.58 (4.03 - 7.71) | 23.9 (21.1 - 26.9) | **3.51 (3.00 - 4.11)** |
| | + choline | 41 | 917 | | 4.3 (3.2 - 5.8) | |
| R2 | -choline | 42 | 1867 | 0.95 (0.61 - 1.47) | 2.2 (1.6 - 3.0) | |
| | + choline | 37 | 1562 | | 2.31 (1.7 - 3.2) | |
| R3 | -choline | 51 | 912 | 1.21 (0.81 - 1.80) | 5.3 (4.1 - 6.9) | |
| | + choline | 42 | 915 | | 4.4 (3.3- 5.9) | |
| R4 | -choline | 105 | 799 | 16.56 (7.75 - 35.40) | 11.6 (9.7 - 13.9) | |
| | + choline | 7 | 991 | | 0.7 (0.3 - 1.4) | |
| R5 | -choline | 141 | 847 | 12.82 (6.98 - 23.53) | 14.3 (12.2 - 16.6) | |
| | + choline | 11 | 977 | | 1.1 (0.6 - 2.0) | |
| R6 | -choline | 93 | 824 | 2.07 (1.47 - 2.90) | 10.1 (8.4 - 12.3) | |
| | + choline | 47 | 910 | | 4.9 (3.7 - 6.5) | |

| Biological Replicate Assays | Condition | Pfs16 +ve | Pfs16 -ve | Rate Ratio (± 95%CI) | GCR (± 95%CI) | Adjusted mean Rate Ratio (± 95%CI) |
|---|---|---|---|---|---|---|
| | | | ***P. falciparum* line 3D7** | | | |
| R1 | -choline | 17 | 965 | 1.22 | 1.7(1.1 - 2.8) | |
| | + choline | 14 | 969 | (0.60 - 2.45) | 1.4 (0.9 - 2.4) | |
| R2 | -choline | 1 | 997 | 0.20 | 0.1 (0.0 - 0.6) | |
| | + choline | 5 | 989 | (0.02- 1.70) | 0.5 (0.2 - 1.2) | |
| R3 | -choline | 6 | 987 | 3.01 | 0.6 (0.3 - 1.3) | |
| | + choline | 2 | 995 | (0.61 - 14.88) | 0.2 (0.0 - 0.7) | |
| R4 | -choline | 34 | 931 | 3.45 | 3.5 (2.5 - 4.9) | |
| | + choline | 10 | 970 | (1.72 - 6.95) | 1.0 (0.6 - 1.9) | **1.50 (1.16 - 1.94)** |
| R5 | -choline | 4 | 940 | 0.82 | 0.4 (0.2 - 1.1) | |
| | + choline | 5 | 961 | (0.22 - 3.04) | 0.5 (0.2 - 1.2) | |
| R6 | -choline | 6 | 987 | 0.65 | 0.6 (0.3 - 1.3) | |
| | + choline | 9 | 961 | (0.23 - 1.82) | 0.9 (0.5 - 1.8) | |
| R7 | -choline | 35 | 931 | 1.86 | 3.6 (2.6 - 5.0) | |
| | + choline | 19 | 961 | (1.08 - 3.24) | 1.9 (1.2 - 3.0) | |
| R8 | -choline | 43 | 913 | 1.28 | 4.5 (3.4 - 6.0) | |
| | + choline | 34 | 938 | (0.83 - 1.99) | 3.5 (2.5 - 4.8) | |
| | | | ***P. falciparum* line T9/96** | | | |
| R1 | -choline | 7 | 985 | 1.00 | 0.7 (0.3 - 1.4) | |
| | + choline | 7 | 985 | (0.35 - 2.84) | 0.7 (0.3 - 1.4) | |
| R2 | -choline | 2 | 995 | 0.57 | 0.2 (0.0 - 0.7) | |
| | + choline | 4 | 1131 | (0.10 - 3.10) | 0.4 (0.1 - 0.9) | |
| R3 | -choline | 2 | 995 | | 0.2 (0.0 - 0.7) | |
| | + choline | 0 | 999 | no estimate | 0 (0) | |
| R4 | -choline | 2 | 934 | 0.26 | 0.2 (0.0 - 0.8) | **0.8 (0.51 - 1.19)** |
| | + choline | 8 | 983 | (0.06 - 1.24) | 0.8 (0.4 - 1.6) | |
| R5 | -choline | 5 | 643 | 0.83 | 0.8 (0.3 - 1.8) | |
| | + choline | 4 | 429 | (0.23 - 3.09) | 0.9 (0.4 - 2.4) | |
| R6 | -choline | 11 | 978 | 0.84 | 1.1 (0.6 - 2.0) | |
| | + choline | 13 | 973 | (0.38 - 1.88) | 1.3 (0.8 - 2.2) | |
| R7 | -choline | 9 | 739 | 0.79 | 1.2 (0.6 - 2.3) | |
| | + choline | 12 | 780 | (0.34 - 1.87) | 1.5 (0.9 -2.6) | |

| Biological Replicate Assays | Condition | Pfs16 +ve | Pfs16 -ve | Rate Ratio (± 95%CI) | GCR (± 95%CI) | Adjusted Risk Ratio (± 95%CI) |
|---|---|---|---|---|---|---|
| | | | ***P.falciparum*** **line NF54** | | | |
| R1 | -choline | 27 | 945 | 2.50 (1.25 - 5.01) | 2.8 (1.9 - 4.0) | **2.41 (1.97 - 2.96)** |
| | + choline | 11 | 978 | | 1.1 (0.6 - 2.0) | |
| R2 | -choline | 30 | 943 | 5.11 (2.13 - 12.24) | 3.1 (2.2- 4.4) | |
| | + choline | 6 | 990 | | 0.6 (0.3 - 1.3) | |
| R3 | -choline | 40 | 819 | 3.53 (1.90 - 6.56) | 4.7 (3.4- 6.3) | |
| | + choline | 13 | 973 | | 1.3 (0.8 - 2.2) | |
| R4 | -choline | 43 | 913 | 5.57 (2.63 - 11.79) | 4.5 (3.4 - 6.0) | |
| | + choline | 8 | 983 | | 0.8 (0.4 - 1.6) | |
| R5 | -choline | 70 | 861 | 1.52 (1.06 - 2.18) | 7.5 (6.0 - 9.4) | |
| | + choline | 47 | 905 | | 4.9 (3.7 - 6.5) | |
| R6 | -choline | 60 | 880 | 1.86 (1.23 - 2.83) | 6.4 (5.0 - 8.1) | |
| | + choline | 33 | 933 | | 3.4 (2.4 - 4.8) | |
| R7 | -choline | 8 | 983 | 1.60 (0.53 - 4.87) | 0.8 (0.4 - 1.6) | |
| | + choline | 5 | 986 | | 0.5 (0.2 - 1.2) | |
| R8 | -choline | 15 | 970 | 5.06 (1.47 - 17.41) | 1.5 (0.9 - 2.5) | |
| | + choline | 3 | 993 | | 0.3 (0.1 - 0.9) | |

**Supplementary Figure S1.** Scheme outlining the timing of processes undertaken for two different methods to assay parasite gametocyte conversion rates of different *P. falciparum* lines in culture (details given in Materials and Methods section).

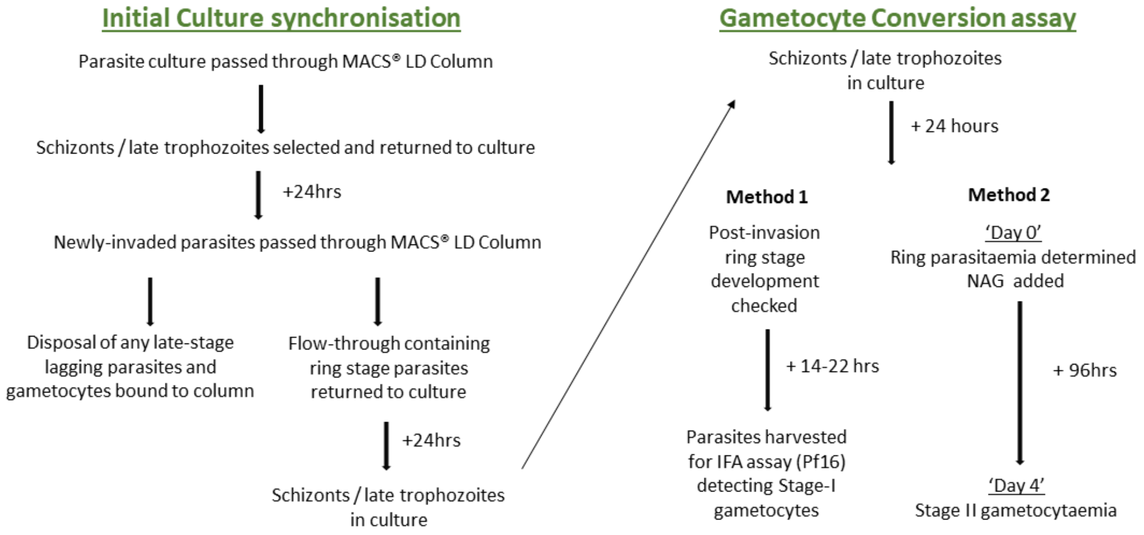
